# Thermophilization, climatic debt, and consequent declines in primary productivity

**DOI:** 10.64898/2026.04.28.721369

**Authors:** Yuna Daido, Kaede Konrai, Yusuke Onoda, Shinichi Tatsumi

## Abstract

Species have optimal environmental conditions, and ongoing climate warming is reshaping community composition. In particular, many ecosystems exhibit thermophilization, a shift toward species adapted to warmer conditions. However, this process is often slower in forests, leading to a mismatch between community composition and ambient temperature, referred to as climatic debt. Despite increasing attention, its effects on forest productivity remain unclear. Quantifying tree community responses to warming is therefore essential for predicting future forest dynamics and informing biodiversity conservation.

In this study, we analyzed natural forests across Japan using data from the 3rd and 4th National Forest Inventory periods (2009–2018). We first assessed compositional consistency between survey periods using the Bray–Curtis index and excluded plots with high dissimilarity (≥ 0.8). Species-specific thermal optima were estimated using species distribution models and used to calculate the Community Temperature Index (CTI). Thermophilization was quantified as the temporal change in CTI, while climatic debt was defined as the difference between CTI and mean annual temperature. We then examined the relationship between climatic debt and changes in aboveground biomass, used as a proxy for productivity, using linear mixed-effects models.

We found a mean thermophilization rate of 0.005 °C yr⁻¹. Despite this shift, climatic debt increased at an average rate of −0.022 °C yr⁻¹, indicating a growing mismatch between climate warming and community thermal composition. Although thermophilization showed no statistically significant association with stand structure, it tended to vary with the proportion of small-diameter trees, suggesting the influence of multiple interacting drivers. Importantly, increasing climatic debt was significantly associated with declines in forest primary productivity, even after accounting for stand structure and regional variation. These results demonstrate that delayed thermal adjustment of tree communities can constrain forest productivity under ongoing climate warming, highlighting the importance of evaluating community-level thermal responses for sustaining forest ecosystem functioning.

## 1. Introduction

Global warming forces species to shift toward cooler latitudes and higher elevations as they track their thermal niches (Woodward & Williams, 1987; Thuiller, 2004). Such distributional shifts can alter composition of local biological communities by increasing the relative abundance of species adapted to warmer climatic conditions. This directional change in community composition is referred to as “thermophilization” (Gottfried et al., 2012). Thermophilization occurs through two complementary demographic processes: the increasing recruitment or dominance of warm-adapted species whose thermal optima exceed those of co-occurring taxa, and the concomitant decline of species adapted to cooler conditions (McLean et al., 2021). Thermophilization is typically quantified by temporal changes in the community temperature index (CTI), which represents the community-weighted mean of species’ temperature affinities (Devictor et al., 2012; Khaliq et al., 2024) (Fig. S1). Increasing CTI values indicates transitions toward communities dominated by warm-adapted species, a pattern that has been documented in ecosystems worldwide (Rosenblad et al., 2023).

Thermophilization often lags behind the pace of contemporary temperature rises (Bertrand et al., 2011; Morin, 2011).The resultant mismatch between CTI and ambient temperature is referred to as climatic debt (Bertrand et al., 2016) . Accumulation of climatic debt has been found across multiple types of ecosystems, among which forests have been one of the most significant, given the decadal to centennial timescales of tree mortality, generating strong demographic inertia (Aitken et al., 2008; Zhu et al., 2012). In addition to the slow demographic processes, dispersal limitations can restrict the colonization of warm-adapted tree species, thereby increasing climatic debt (Alexander et al., 2017). Biotic interactions, including competitive effects and microclimatic buffering by existing canopy trees on newly colonized trees, can further delay thermophilization (De Frenne et al., 2013). These ecological characteristics of forest ecosystems make them a suitable model system for understanding the ecological mechanisms underlying delayed biotic responses to climate warming (Bertrand et al., 2016; Rosenblad et al., 2023).

Global warming can reduce forest primary productivity by inducing physiological stress (Hogan et al., 2024). Such stresses are expected to be particularly pronounced in forests with high climatic debt—that is, tree communities dominated by species with thermal niches aligned with past, cooler climates and/or those lacking recruitment of species adapted to current thermal conditions. These mismatches between species thermal composition and ambient temperature can thus play a key role in climate-induced declines in forest productivity (Cuni-Sanchez et al., 2024). However, empirical evidence linking thermophilization, climatic debt, and forest productivity remains scarce. Filling the gap among these linkages is essential for understanding forest ecosystem functioning under climate change, as well as improving the accuracy of global vegetation projections, which often assume an instant or no responses of plant communities to warming.

Here, we quantified thermophilization and climatic debt in natural forests, analyzed the consequences of climatic debt magnitude on primary productivity. We hypothesized that: (1) While thermophilization is occurring, its pace lags behind the rate of temperature warming, leading to the accumulation of climate debt, and (2) Forests experiencing larger climatic debt exhibit lower productivity. To test these hypotheses, we used the National Forest Inventory (NFI) dataset in Japan. Despite the ecological importance of East and Southeast Asian countries that harbor high biodiversity, there is little study on thermophilization and climatic debt in these region compared to European or North American countries (De Frenne et al., 2013; Feeley et al., 2020). Our study provides the first national-scale empirical assessment linking thermophilization, climatic debt, and primary productivity in natural forests in one of the world’s biodiversity hotspots.

## 2. Materials and Methods

All data processing and analyses were conducted using R version 4.4.3 (R Core Team., 2025).

### 2.1 Plots and Species Data

We used NFI dataset collected by the Forestry Agency of Japan (Forest Agency., 2017). We analyzed the third (2008–2013) and fourth (2013–2018) surveys. From all NFI plots, we retained only those classified as natural forests in both cycles. We further limited the dataset to continued plots. Although some plots were spatially mismatched between the two time periods (Nakajima., 2017), this study used only plots with consistent locations across both periods (see the Appendix 10.1 and 10.2 for details on data filtering). The final dataset used in the subsequent analyses comprised 4,412 plots.

### 2.2 Climate Data

We used the climate mesh dataset (2012) and annual precipitation data (2012) from the Geospatial Information Authority of Japan (Ministry of Land, Infrastructure, Transport and Tourism., 2012). We also used daily mean temperature mesh data from 1990 to 2020, published by the National Agriculture and Food Research Organization (NARO), to derive more detailed temperature variables (NARO., 2022).

### 2.3 Estimation of Species’ Optimal Temperature (DT_opt_)

We estimated the optimal temperature of distribution (DT_opt_) for each species to calculate CTI, based on presence–absence data from the third survey and climate data. We defined DT_opt_ as the temperature at which the modeled probability of occurrence reached its maximum. We excluded six plots because of data deficiency. For species affected by taxonomic inconsistencies, we used revised species labels and merged presence–absence data for the paired taxa accordingly.

We conducted DT_opt_ estimation in two steps (Fig. S4). First, we separated species based on the frequency of presence records. We used a threshold of 50 presence observations because this number was the minimum number required to allow all models in the subsequent modeling procedure to run without errors. We modeled species with more than 50 presence records (n = 202) using the Biomod2 package in R, whereas we assigned DT_opt_ values to species with fewer than 50 presence records (n = 673) based on the mean annual temperature of plots where we recorded the species.

For species exceeding the 50-record threshold, we fitted seven single models (ANN, CTA, FDA, GLM, MARS, SRE, and XGBoost) and repeated each model 10 times. We evaluated model performance using the True Skill Statistic (TSS). We constructed ensemble models using four approaches: the ensemble means, ensemble median, committee averaging, and weighted mean ensemble. We used the weighted mean ensemble for subsequent analyses. We repeated this procedure for all species that met the more-than-50-observation criterion.

### 2.4 Calculation of Community Temperature Index (CTI) and Thermophilization

We calculated CTI (°C) for each plot using the DT_opt_ values obtained in the previous step and the biomass of each species present in the plot, according to the following equation:

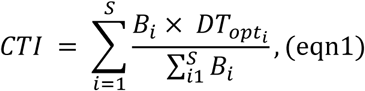

where *B_i_* is total biomass of species i, *DT_opti_* is optimal temperature of species i, and *S* is the number of total species in the plot.

We calculated CTI separately for each NFI cycle. We then defined thermophilization (°C yr⁻¹) as the temporal change in CTI between the two survey periods.

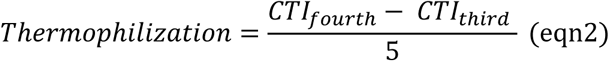

### 2.5 Analysis of thermophilization

We also analyzed the underlying mechanism of thermophilization. We first evaluated the relationship between thermophilization and stand structure using a model that accounted for conifer and evergreen proportions, as well as the relative abundance of small-diameter trees (DBH ≤ 15 cm).

Then, we calculated the individual contribution of each species to thermophilization (contribution), with the following formula (Borderieux et al., 2024a):

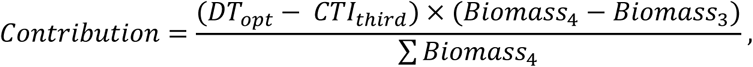

where DT_opt_ is the optimal temperature of species i, Biomass_3_ and Biomass_4_ are the above biomass of species i in third and fourth survey respectively, and ∑ 𝐵𝑖𝑜𝑚𝑎𝑠𝑠_4_ is the total above biomass of the plot. We calculated plot-level contribution values by summing the contributions of all species within each plot.

### 2.6 Estimation of Climatic debt

We then calculated climatic debt. We used the geographic coordinates of all plots to extract local temperature data from the 1990–2020 daily mean temperature dataset provided by NARO. We calculated annual mean temperatures for each year from the daily mean temperature data using the R package supplied by NARO. We fitted a regression curve to the resulting 30-year time series of annual mean temperatures using a generalized additive mixed model (GAMM), which allowed us to explicitly account for temporal autocorrelation and improved the robustness of the estimated temperature trend. We extracted predicted values from the fitted regression curve corresponding to the survey years of the third and fourth NFI cycles and treated them as the annual mean temperatures. We calculated climatic debt (°C yr⁻¹) as follows.

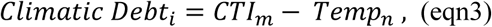

where CTI_m_ is CTI (m is the period when the CTI was calculated) and Temp_n_ is the fitted value on the regression line for year n (the mean temperature of plot i in year n).

### 2.7 Relationship between climatic debt and primary productivity

We quantified plot-level changes in aboveground biomass (AGB) using data from the third survey to evaluate the relationship between climatic debt and forest primary productivity. We defined forest productivity as the relative rate of change in AGB between consecutive census periods. This approach aligns with the IPCC inventory guidelines, in which researchers use changes in aboveground biomass as high-tier indicators for estimating forest carbon sequestration (IPCC, 2006 ; IPCC, 2019).

We employed linear mixed-effects models to analyze drivers of productivity and treated climatic debt and structural variables as fixed effects. We included “Region” as a random intercept to account for spatial autocorrelation and regional environmental heterogeneity. We initially constructed a global model that contained all structural covariates and then performed model selection based on Akaike’s Information Criterion (AIC) to identify the most parsimonious model. We then visualized the effect of climatic debt on productivity using partial effects plots while controlling other covariates. We specified the final model as follows:

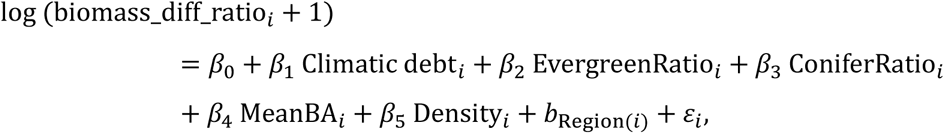

where 𝑏_Region(𝑖)_ represents the random intercept for region, assumed to follow a normal distribution with mean zero, and 𝜀_𝑖_ denotes the residual error.

## 3. Results

### 3.1 Thermophilization

We found that Thermophilization was observed at the national scale. A one-sided test detected a statistically significant positive trend across Japan. The national mean thermophilization rate was 0.026℃ 5yr⁻¹ or 0.005 °C yr⁻¹ (Table 1). The standard deviation was large relative to the mean (SD = 0.35), indicating substantial variability among plots. Among the 4,406 plots analyzed, thermophilization values were positive in more than half of the plots.

**Table 1.**
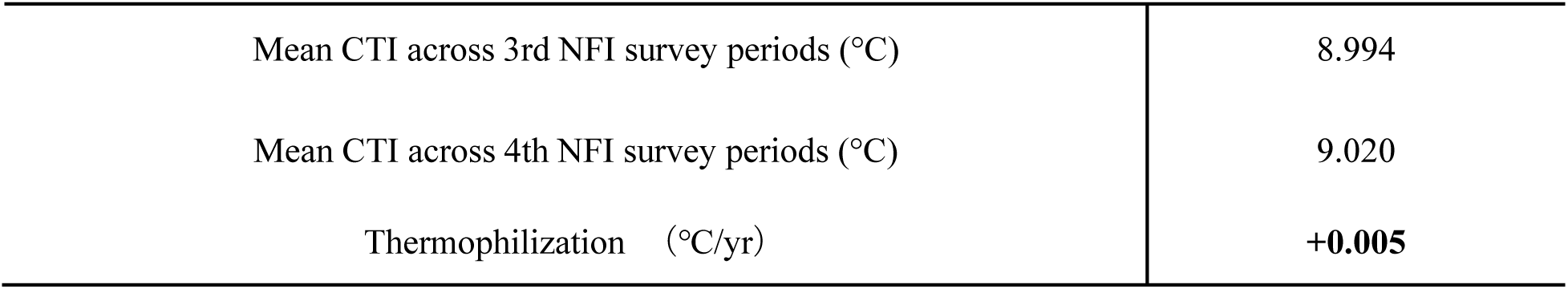
Thermophilization across survey periods.

When plots were grouped by forestry regions defined by the Forestry Agency of Japan, thermophilization differed significantly among regions (Welch’s ANOVA, *F* = 2.48, *p* = 0.024). Since the sample size in Okinawa was relatively small (n = 21), a sensitivity analysis excluding this region was conducted, and the regional differences remained significant (*F* = 2.98, *p* = 0.010), indicating that the results were robust to its inclusion (Fig.1, Table S2).

**Figure 1.**
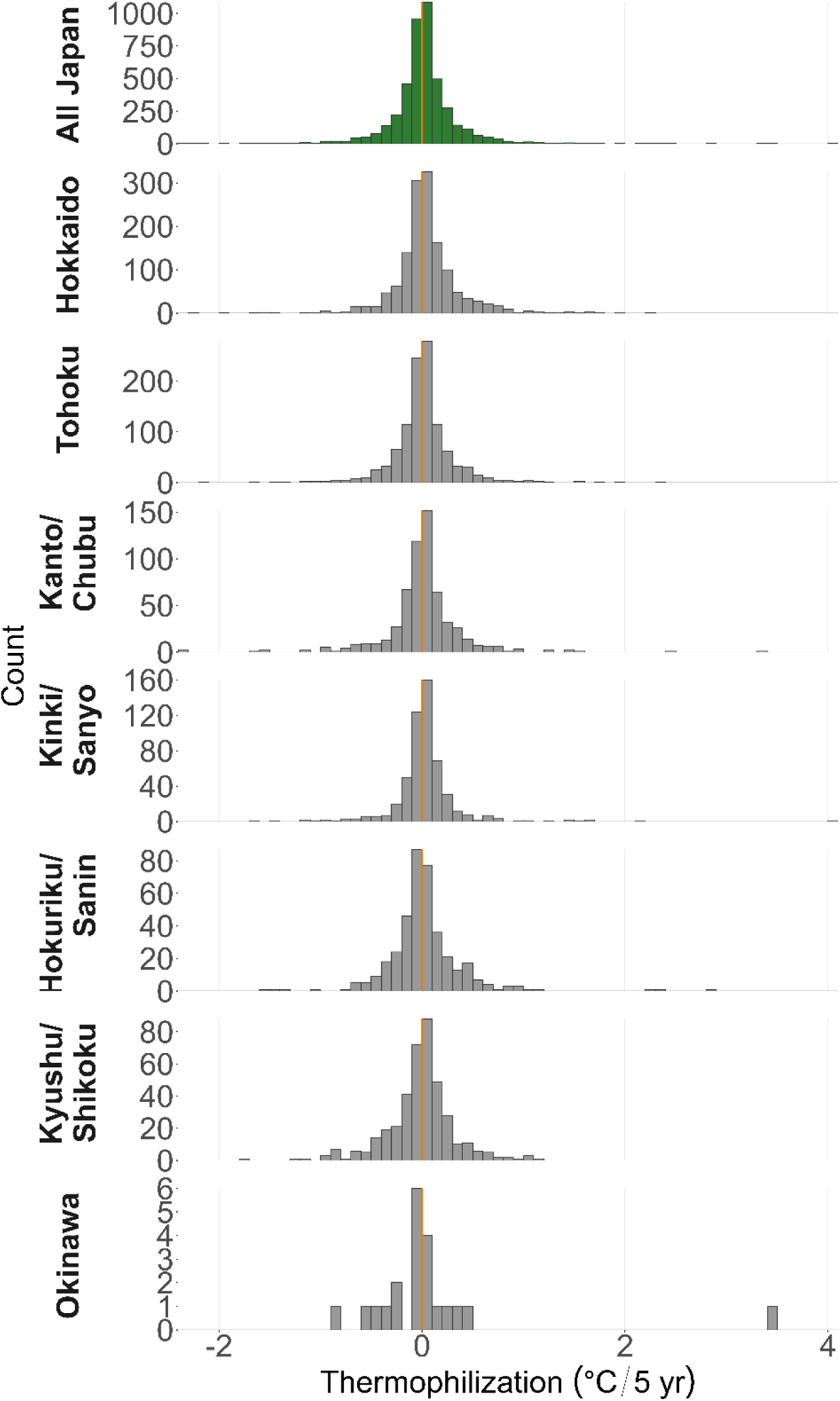
Distribution of thermophilization across Japan and major regions defined by the Forestry Agency of Japan. The green histogram represents the nationwide distribution, whereas gray histograms show region-specific distributions. The dashed orange vertical line indicates zero thermophilization.

At the plot level, clear differences in contribution patterns were observed depending on the direction of thermophilization. In plots where thermophilization progressed, increases in biomass of warm-adapted species and declines in biomass of cold-adapted species showed large contributions, and both were significantly associated with the thermophilization index (Fig. 2). In contrast, in plots where thermophilization showed negative changes, decreases in biomass of warm-adapted species and increases in biomass of cold-adapted species made relatively larger contributions. Although their effects were weaker than those observed in plots with progressing thermophilization, these processes acted in directions that suppressed or reversed thermophilization.

**Figure 2.**
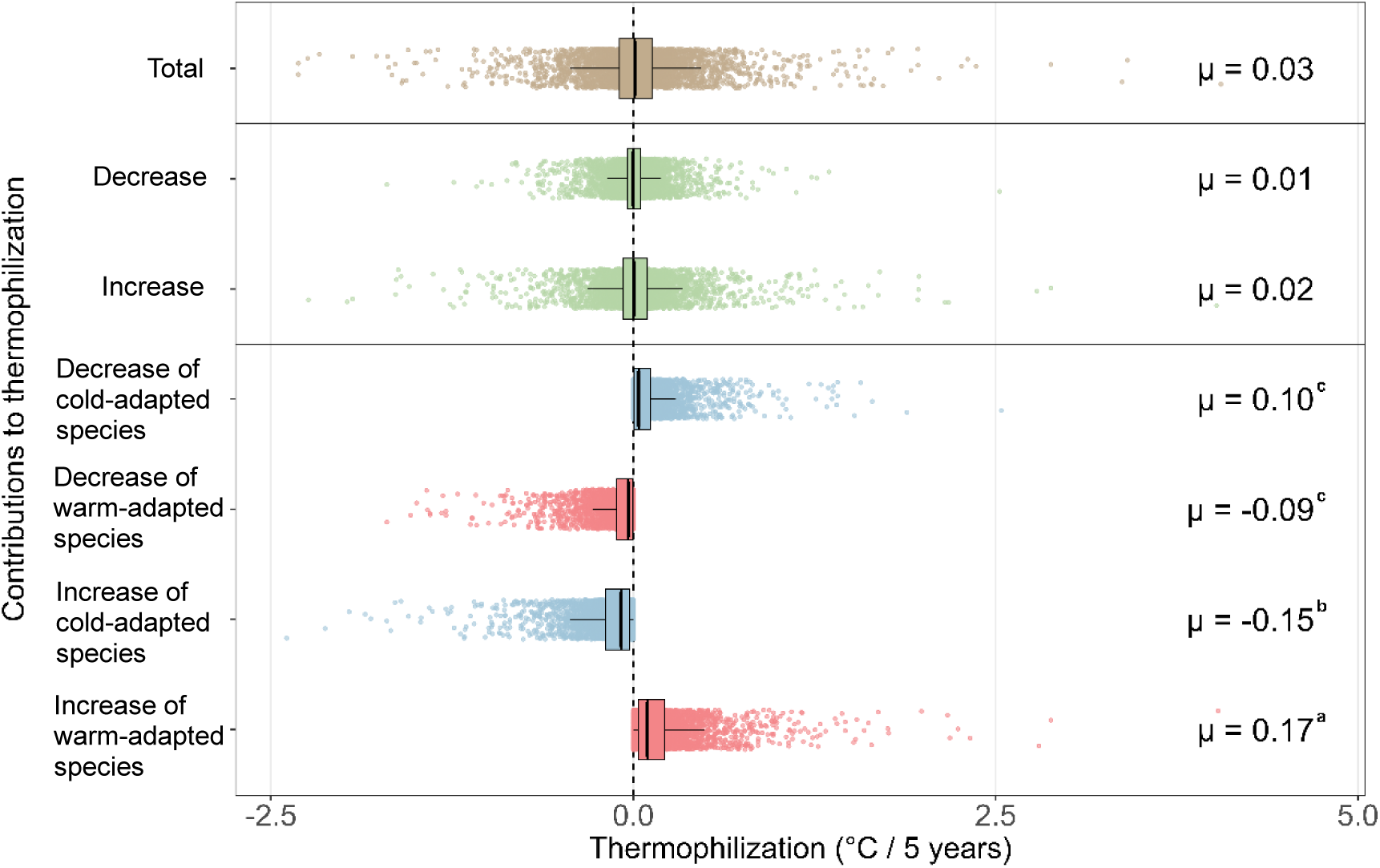
Contributions to thermophilization across demographic processes. Distributions of plot-level contributions are shown for the total community, decrease, increase, and four demographic components classified by thermal affinity (cold-vs. warm-adapted species). Boxes show medians and interquartile ranges; points represent individual plots. Mean values (μ) are indicated on the right. Letters represent significant differences among the four demographic components based on ANOVA of absolute contributions with Tukey’s HSD test.

In further analysis that included variables for stand structure, we adopted model averaging because the AIC values of the compared models showed little to no difference (Table 2). Based on this approach, the proportion of conifers, the proportion of evergreen trees, and the proportion of small trees were all found to be non-significant. Furthermore, the variance of thermophilization increased significantly with an increasing proportion of trees with a small diameter or conifer trees (Fig. S6).

**Table 2.** Model selection based on AIC. The candidate models are shown and ranked by AICc. Fixed effects include climate variables and stand structural variables, with region included as a random intercept. The top six models were used for model averaging.

| **Small trees** | **Evergreen** | **Conifer** | **df** | **AICc** | **ΔAICc** | **AIC weight** |
| --- | --- | --- | --- | --- | --- | --- |
| -0.051 | NA | -0.032 | 5 | 3328.5 | 0 | 0.222 |
| -0.042 | -0.026 | NA | 5 | 3328.87 | 0.37 | 0.185 |
| -0.039 | NA | NA | 4 | 3329.25 | 0.76 | 0.152 |
| NA | NA | NA | 3 | 3329.76 | 1.26 | 0.118 |
| NA | -0.023 | NA | 4 | 3329.92 | 1.42 | 0.109 |
| -0.049 | -0.012 | -0.022 | 6 | 3330.29 | 1.79 | 0.091 |
| NA | NA | -0.021 | 4 | 3330.5 | 2 | 0.082 |
| NA | -0.02 | -0.004 | 5 | 3331.9 | 3.4 | 0.041 |

### 3.2 Climatic debt

A statistically significant change in climatic debt was also detected across Japan. The mean climatic debt was −0.316 °C in the 3rd NFI survey and −0.427 °C in the 4th survey. The average rate of change in climatic debt was −0.022 °C yr⁻¹ over the five-year (Table 3, Fig. 3).

**Table 3.** Climatic debt across survey periods.

|  |  |
| --- | --- |
| Climatic debt (3rd NFI period, °C) | -0.316 |
| Climatic debt (4th NFI period, °C) | -0.427 |
| Rate of climatic debt (°C/yr) | **-0.022** |

**Figure 3.**
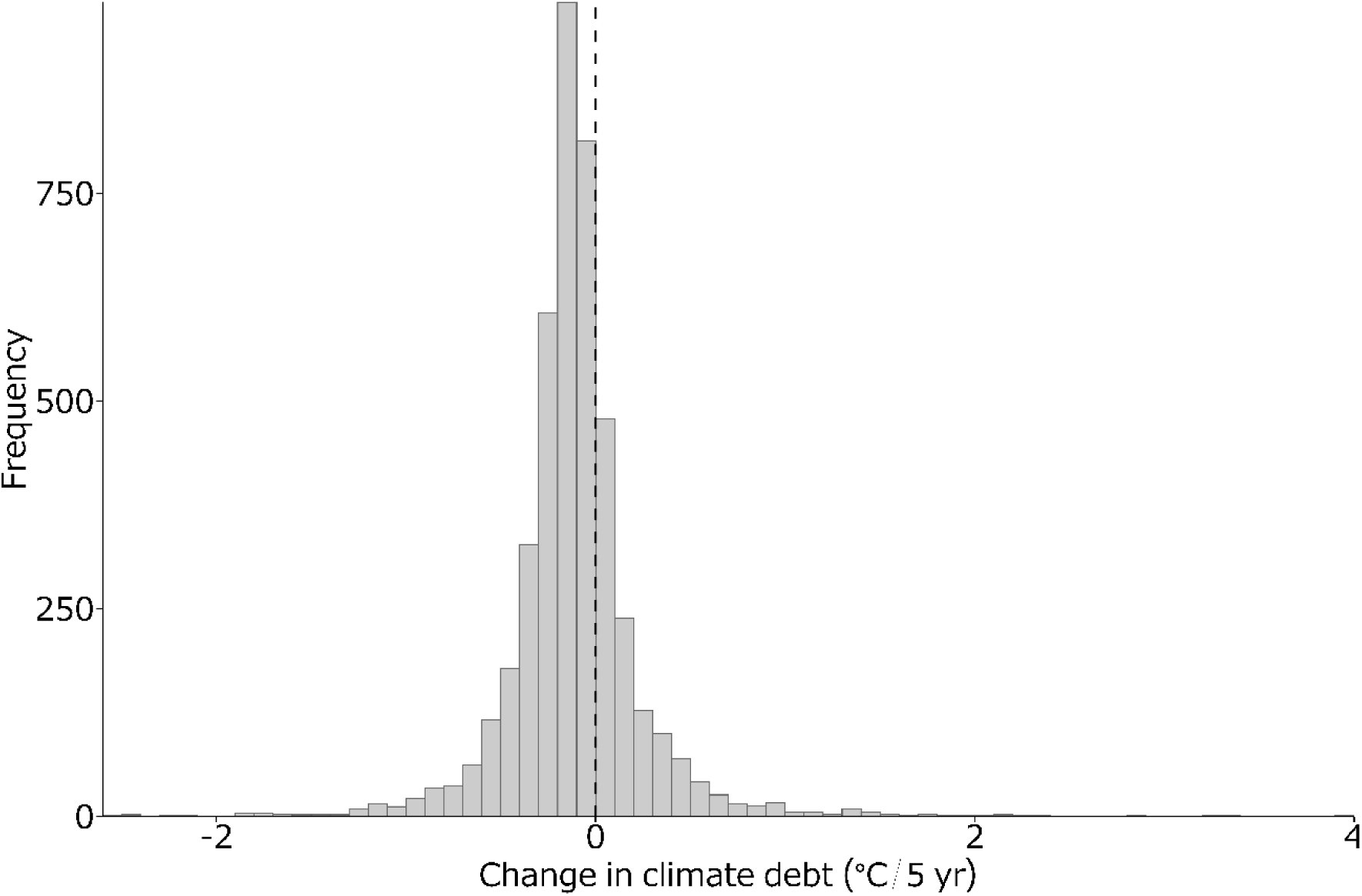
Changes in climatic debt. Histogram shows the change in climatic debt between the third and fourth survey periods. Negative values indicate increasing climatic debt, whereas positive values indicate a reduction. The dashed vertical line denotes zero change.

As defined in Eqn3, positive change of climatic debt (CTI_fourth_ – CTI_third_ > 0) indicates a reduction in climatic debt, whereas negative change (CTI_fourth_ – CTI_third_ < 0) indicates an increase in climatic debt. The observed negative mean change therefore indicates an increase in climatic debt between the third and fourth NFI surveys.

### 3.3 Productivity

At the national scale, the linear mixed-effects model revealed a significant relationship between climatic debt and forest productivity (Table 4). We found that productivity declined as climatic debt increased. Since more negative climatic debt values represent a larger climatic lag of tree communities relative to temperature increases, this result indicates that plots experiencing greater climatic debt exhibited lower forest productivity (Fig. 4).

**Figure 4.**
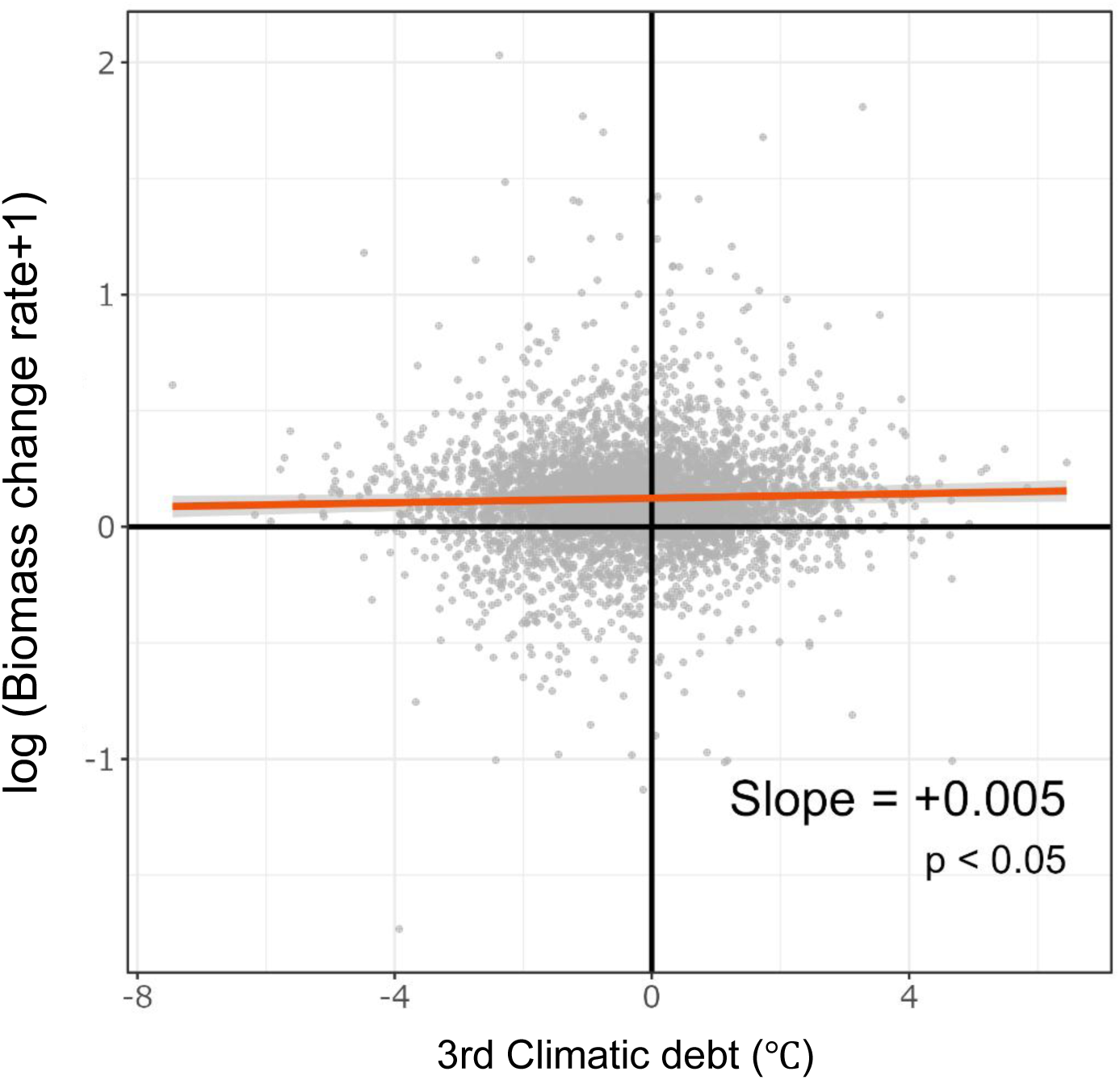
Relationship between forest productivity and climatic debt. The figure shows the relationship between forest productivity and climatic debt. Values on the x-axis represent climatic debt, with more negative values indicating a greater progression of climatic debt.

**Table 4.** Model selection based on AIC. Model selection results for linear mixed-effects models explaining variation in biomass difference ratio. The four candidate models with non-zero Akaike weights are shown and ranked by AICc. Fixed effects include evergreen ratio, conifer ratio, mean basal area, and stem density, with region as a random intercept. The top-ranked model (lowest AICc) was selected for subsequent analyses.

| Evergreen ratio | Conifer ratio | Mean basal area | Stem density | df | AICc | ΔAICc | AIC weight |
| --- | --- | --- | --- | --- | --- | --- | --- |
| 0.124 | -0.102 | -3.342 | -0.013 | 7 | 3330.467 | 0 | 0.982 |
| 0.043 | NA | -3.374 | -0.012 | 6 | 3338.963 | 8.495 | 0.014 |
| NA | NA | -3.258 | -0.012 | 5 | 3342.355 | 11.888 | 0.003 |
| NA | -0.001 | -3.254 | -0.012 | 6 | 3344.355 | 13.887 | 0.001 |

After log-transforming the biomass change rate to satisfy model assumptions, climatic debt remained a significant predictor even after accounting for stand structural attributes, including evergreen ratio, conifer ratio, mean basal area, and tree density, as well as regional differences modeled as a random intercept.

## 4. Discussion

### 4.1 Thermophilization progression and climatic debt expansion

We found a statistically significant thermophilization and climatic debt, supporting our first hypothesis (Table1 and Table 3). This trend was consistent across regions, suggesting a coherent response of tree communities to ongoing climate warming. As for thermophilization, such a pattern is in line with earlier findings and implies that warming has altered community composition through the relative increase and differential growth of species with higher thermal optima (Gottfried et al., 2012). The estimated rate of thermophilization, which was 0.005 °C yr⁻¹, falls within the range reported for other temperate regions, indicating that the magnitude observed in this study is ecologically plausible (Fadrique et al., 2018 ; Chen et al., 2025; Cuni-Sanchez et al., 2024; Khaliq et al., 2024; Rosenblad et al., 2023). At the same time, our results show that climatic debt has deepened across natural forests, indicating an increasing mismatch between community thermal composition and ambient temperature (Table 2, Fig. 3). This pattern suggests that tree communities have not kept pace with the rapid rate of recent climate warming although thermophilization occurs. Similar delays in community-level responses have been widely reported in forest ecosystems, particularly natural forests (Bertrand et al., 2011; Borderieux et al., 2024b; Richard et al., 2021; Rosenblad et al., 2023) (Khaliq et al., 2024; Rosenblad et al., 2023).

This mismatch might be explained by examining species turnover (Fig. 2). The increase in biomass of warm-adapted species (μ = 0.17) exerted the strongest influence and was greater than the decrease in cold-adapted species (μ = −0.10). Across many forests spanning low- to high-latitude regions, previous studies have reported that declines in cold-adapted species contribute most to thermophilization, whereas increases in warm-adapted species are generally secondary and delayed (Borderieux et al., 2024b; Cuni-Sanchez et al., 2024; Talluto et al., 2017). In contrast, our results indicate that increases in warm-adapted species biomass play the dominant role. This suggests that warm-adapted species are actively increasing in dominance within these forests, rather than thermophilization being driven primarily by the loss of cold-adapted species. Furthermore, plots showing limited thermophilization tended to exhibit larger climatic debt. This pattern implies that climatic debt may emerge in forests where species turnover has not sufficiently tracked recent warming, or where declines in cold-adapted species occur without a compensatory increase in warm-adapted species (McNichol & Russo, 2023). In such cases, community composition lags behind climate change, leading to an expansion of climatic debt (Bertrand et al., 2016).

The delayed decline or even continued increase of cold-adapted species under ongoing warming can be explained by several ecological mechanisms that constrain rapid community restructuring. In natural forests, community composition is strongly influenced by large, long-lived trees that can persist for decades or even centuries, creating substantial demographic inertia (Mäkinen et al., 2025). Such longevity slows compositional turnover and decouples short-term climatic change from immediate shifts in dominance. Even when warm-adapted species are present, increases in their relative biomass depend on slow processes, including recruitment, competitive displacement, and gap-phase dynamics, resulting in delayed thermal realignment at the community level (HilleRisLambers et al, 2015). Dispersal limitation can also restrict the recruitment and establishment of warm-adapted species, while canopy-mediated microclimatic buffering may locally dampen macroclimatic warming, maintaining cooler understory conditions suitable for cold-adapted taxa (Primack & Miao, 1992; Bertrand et al., 2011; De Frenne et al., 2013). These processes cause cold-adapted species to persist despite rising temperatures, thereby generating a mismatch between community composition and contemporary climate.

### 4.2 Variation of thermophilization

In the analysis of stand structure, specific structural variables did not fully explain the mean thermophilization. Although thermophilization was consistently detected across forest stands, no single structural variable showed a strong and unambiguous effect. Based on the sum of weights in model averaging, the proportion of small trees exhibited the highest relative importance (0.74) and the most negative regression coefficient (Fig. S5). However, this effect was accompanied by substantial uncertainty, as the 95% confidence intervals included zero. These results suggest that while stand structure may exert a secondary influence on the average thermophilization signal, the observed national-scale trend is more likely driven by the combined effects of microclimate conditions, site characteristics, and past disturbance history rather than stand structure alone (De Frenne et al., 2013; Kiebacher et al., 2023; Rosenblad et al., 2023). Therefore, interpretations of structural effects on mean thermophilization should be made cautiously.

In contrast to the weak structural control on the mean response, stand structure was closely associated with the variability of thermophilization. The variance increased with the proportion of small or conifer trees, and thermophilization values exhibited both positive and negative changes (Fig. S6). Such pronounced variability may reflect differences in physiological traits, regeneration strategies, and heterogeneous responses to local environmental conditions (De Frenne et al., 2013; Kiebacher et al., 2023). Importantly, this variability does not necessarily indicate rapid or unidirectional adaptation to warming but rather suggests the existence of diverse response pathways within forest communities (Fadrique, 2018).

### 4.3 Climatic debt and productivity

Our results demonstrate that forest productivity significantly declined as climatic debt increased, providing empirical support for our second hypothesis (Fig. 4, Table 4). Even after accounting for regional heterogeneity and stand structural attributes such as basal area, tree density, and functional composition, plots with more negative climatic debt values consistently exhibited lower productivity. This finding suggests that climatic debt is not merely a localized phenomenon or a structural factor. Instead, it represents a pervasive constraint on forest functioning across the diverse environmental conditions of Japanese natural forests (Hogan et al., 2024).

One plausible explanation for this relationship is that in areas where thermophilization has not sufficiently progressed, tree communities remain dominated by species whose optimal temperature are lower than current temperature conditions, which is referred to climatic debt (Hogan et al., 2024; Rosenblad et al., 2023) . Such thermal mismatches may reduce photosynthetic efficiency, increase respiratory costs, and impose physiological stress, ultimately limiting biomass accumulation (Crous et al., 2022; Das et al., 2023; Reich et al., 2016). In addition, rising temperatures are often accompanied by intensified drought stress, which can further constrain tree growth and carbon gain (Gampe et al., 2021; Liu et al., 2021). These mechanisms provide a biologically plausible link between delayed community-level thermal adjustment and reduced forest productivity (Feeley et al., 2020).

Some previous studies have examined warming effects on forest productivity through species-specific physiological responses (Matula et al., 2023). While these approaches have provided valuable insights into how individual species respond to rising temperatures, they may overlook the community-level processes that ultimately shape ecosystem functioning. In contrast, our study demonstrates that climatic debt, defined at the community scale, is significantly associated with changes in forest productivity. By integrating shifts in species composition with productivity responses, our approach moves beyond single-species perspectives and captures ecosystem-level consequences of delayed community adjustment to climate warming. Importantly, this framework incorporates multiple mechanisms simultaneously, including species replacement, growth advantages of warm-adapted species, and environmental constraints such as heat and drought stress. As a result, it provides a more comprehensive understanding of how climatic debt emerges and how it constrains forest productivity undergoing climate change.

However, these findings should be interpreted within the context of the relatively short five-year temporal window examined in this study. Over such a period, stochastic processes, exemplified by individual growth fluctuations and localized mortality, may partially obscure deterministic responses to warming (Chen et al., 2025). While our results highlight that climatic debt can have a negative impact on productivity in the short term, longer-term dynamics may involve compensatory shifts in species composition or functional trait plasticity. Future studies utilizing longer time series will be essential to clarify the multi-decadal interactions between climatic debt and forest carbon dynamics.

Overall, our findings suggest that climatic debt represents a critical constraint on forest productivity under climate warming, emphasizing the importance of community-level thermal adjustment for maintaining ecosystem functioning. By linking thermophilization dynamics with productivity outcomes, this study provides empirical evidence that delays in biotic responses to climate change can have tangible consequences for forest carbon dynamics.

### 4.4 Future Perspectives

While this study provides novel insights into the impacts of climate warming on Japanese forests, it also delineates clear directions for future empirical advancement. Importantly, the present findings establish a quantitative baseline for understanding thermophilization, climatic debt, and their productivity consequences in natural forest ecosystems. Moving forward, several avenues can further deepen and refine these insights.

First, complete plot-level spatial consistency between NFI survey periods could not be fully ensured. Minor spatial mismatches and potential species misidentification may have introduced additional uncertainty into estimates of thermophilization and biomass change. Nonetheless, by applying strict screening criteria and retaining only spatially reliable plots, this study provides a robust analytical framework upon which future refinements can build. Continued improvements in plot relocation accuracy and taxonomic consistency will further enhance the precision of thermophilization assessments.

Second, while the present analysis focused on plot-level patterns, this scale was sufficient to reveal emergent links between climatic debt and forest productivity. Moving forward, analyses incorporating individually tracked trees across survey cycles would allow direct evaluation of demographic processes such as mortality, growth, and recruitment, thereby enabling stronger causal inference regarding the mechanisms underlying delayed community responses to warming.

Third, this study was based on a relatively short observation window of approximately five years, encompassing two consecutive inventory cycles. Despite this limitation, the detection of consistent thermophilization signals and productivity responses over such a short period underscores the sensitivity of Japanese forests to ongoing climate warming. The integration of newly released and future NFI data will facilitate longer-term analyses, providing deeper insights into the temporal dynamics of thermophilization, climatic debt, and forest productivity.

## 5. Conclusion

We found clear evidence of nationwide thermophilization in natural forests, indicating a common shift in tree community thermal characteristics under ongoing climate warming. In addition, climatic debt increased over time and was significantly associated with reduced forest productivity, even after accounting for stand structure and regional differences. This result demonstrates that mismatches between community thermal affinities and ambient temperatures have measurable functional consequences for forest ecosystems.

These findings highlight that evaluating the impacts of climate warming on forest productivity requires consideration of ecological processes such as species dispersal limitation and biotic interactions. Furthermore, our results demonstrate the importance of community-level analyses when assessing ecosystem responses to climate change.

## 6. Data availability

Forest inventory data were obtained from the National Forest Inventory (http://forestbio.jp/). Climate data were sourced from the National Land Numerical Information download service provided by Geographical Survey Institute (https://nlftp.mlit.go.jp/ksj/gml/datalist/KsjTmplt-G02.html) and National Agriculture and Food Research Organization (https://www.naro.go.jp/english/).

The R code and processed datasets used in this study can be downloaded from GitHub (https://github.com/Harmony531/thermophilizationjapan.git).

## Acknowledgements

I am grateful to Dr. Fumio Kitahara for providing the well-organized NFI biomass data used in this study. This work was supported by JSPS KAKENHI Grant Number 24H01527.

## 10. Appendix

### 10.1 Data Selection

Firstly, when we selected plots, we chose “continuing plot” which we defined as sites that met the following criteria: (1) plots that we carried over from the 2nd to the 3rd cycle and that continued into the 4th cycle, or (2) plots that we newly established in the 3rd cycle and that continued into the 4th cycle. In total, 4,780 plots met these criteria. Among them, 8 plots (PlotID: 10448, 10952, 11789, 12422, 12767, 30128, 60004, 320399) lacked tree measurement records in both cycles. We excluded them because we were not able to assess community turnover at these sites. Additionally, several plots contained vegetation records in only one of the two surveys (5 in the 3rd NFI and 3 in the 4th). We removed these plots for the same reason, which resulted in an initial dataset of 4,764 plots.

### 10.2 Data filtering

Before conducting the main analysis, we assessed the consistency of the NFI plot data. Previous research has reported that NFI plot locations do not always correspond across survey cycles, even for plots labeled as “continued,” which suggests potential shifts in actual survey locations over time. Since this study aims to detect temporal changes in species distribution, we ensured spatial correspondence of plots across survey cycles by evaluating the degree of mismatch between the third and fourth surveys.

For each of the 4,764 target plots, we quantified community similarity between the two survey cycles using the Bray–Curtis dissimilarity index, which we calculated from tree species composition and their basal area per plot. We also visually inspected species-by-DBH distributions for each plot. We found several instances of apparent taxonomic misidentification between cycles, which may have led to misdirected species regeneration (Fig. S2). We systematically identified problematic taxonomic replacements by tabulating all species pairs that exhibited such turnover, and we observed that frequent replacements were particularly pronounced within the genus *Acer*. We examined in detail species pairs that switched identities in more than 20 plots. We excluded species pairs that we interpreted as reflecting actual demographic processes rather than taxonomic confusion. For the remaining combinations that we considered likely to represent misidentification, we assigned a unified species label and used it in all subsequent analyses (Table S1). We selected a threshold of 21 plots because replacements occurring fewer times were accompanied by clear increases in the abundance of the new species, which is consistent with natural regeneration rather than misidentification. We assumed that plots with extremely high dissimilarity values likely reflected spatial mismatches or major disturbances. Therefore, we excluded plots with Bray–Curtis values greater than 0.8 (n = 343) from the study (Fig. S3).

**Figure S1.**
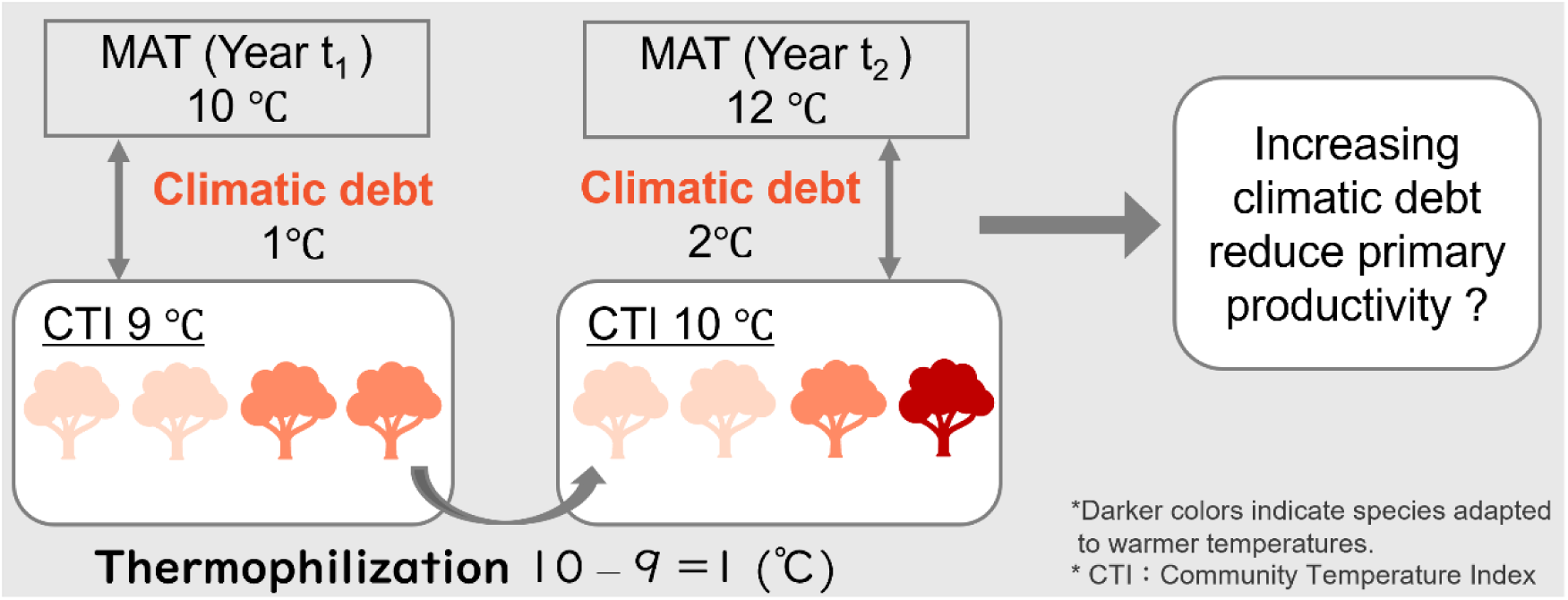
Conceptual illustration of thermophilization, climatic debt, and our hypnoses. Rising temperatures drive shifts in tree community composition toward species associated with warmer climates, a process known as thermophilization. This process is commonly quantified by temporal changes in the Community Temperature Index (CTI), which represents the community-weighted mean thermal affinity of species. Climatic debt arises when increases in CTI lag behind concurrent warming, indicating a mismatch between community thermal composition and ambient temperature.

**Figure S2.**
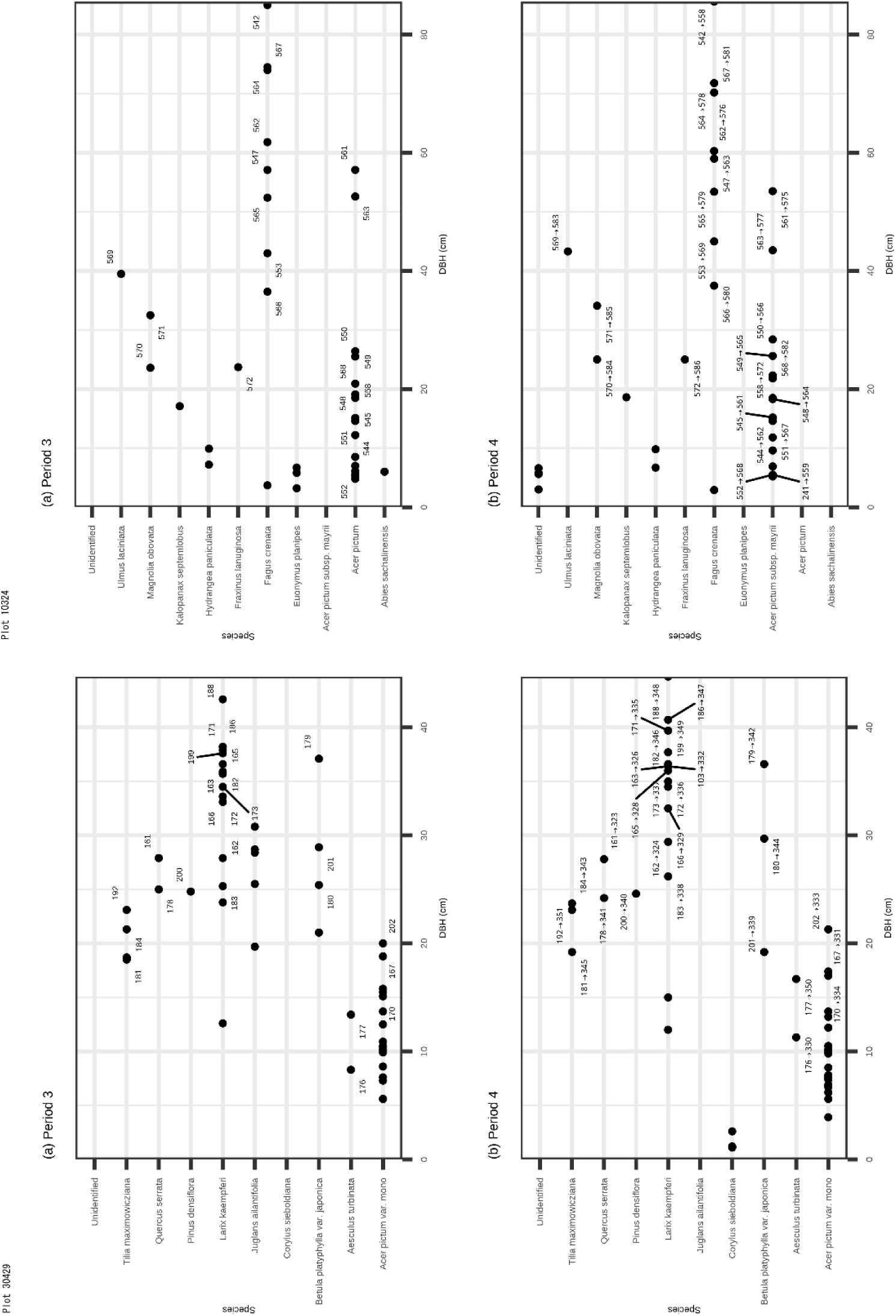
Illustration of tree species turnover between NFI survey periods. The figure shows the correspondence of tree species recorded in the 3^rd^ and 4^th^ NFI survey. Small numbers indicate the identification tags assigned at the time of each survey. In Plot 30429, no species turnover occurred, whereas in Plot 10324, individuals recorded as *Acer pictum* in the 3^rd^ period were recorded as *Acer pictum subsp. mayrii* in the 4^th^ period, illustrating apparent species turnover.

**Figure S3.**
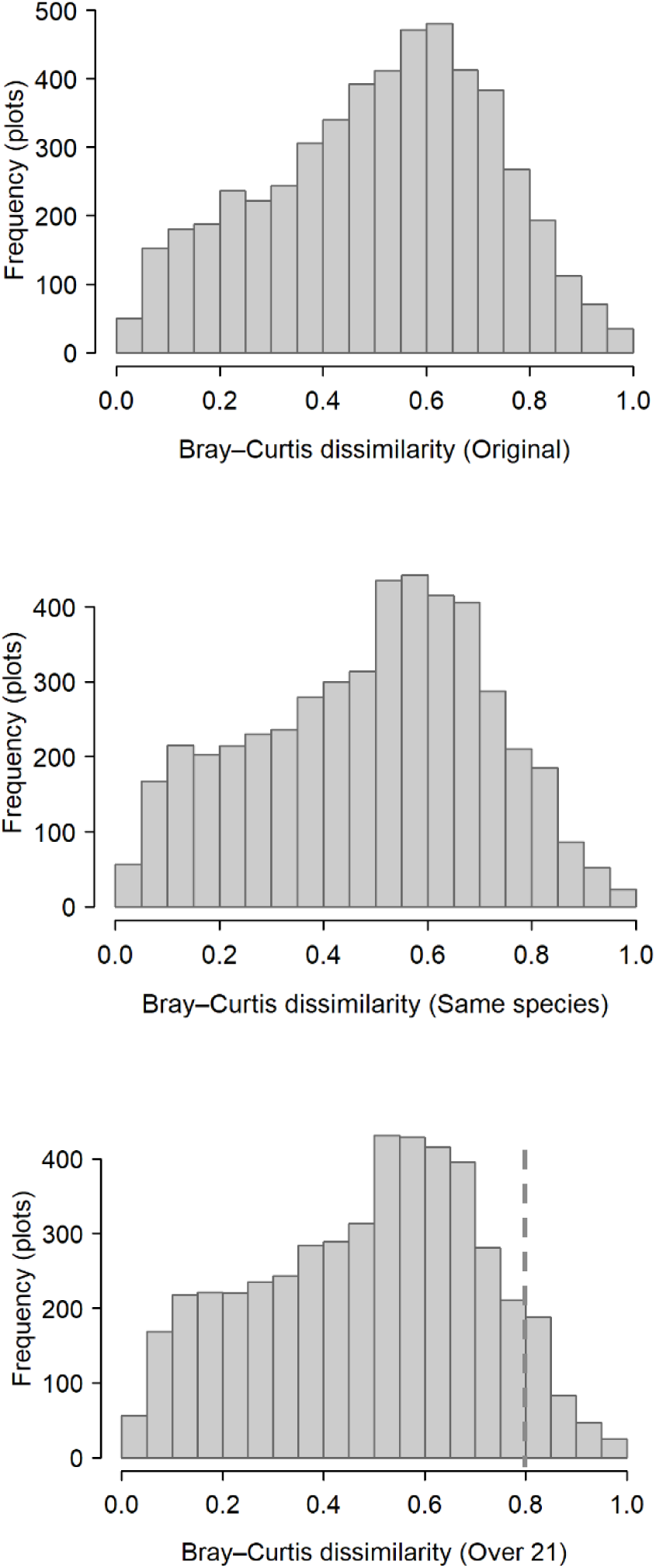
Comparison of Bray–Curtis dissimilarity under different species definitions. Bray–Curtis dissimilarity values were calculated under three data treatments: **Original**, using unmodified species records; **Same Species**, with species names unified only for identical taxa across survey periods; and **Over21**, applying the species definitions used in this study for species exhibiting turnover in at least 21 plots. In this study, we used **Over21** version and its plots with Bray–Curtis dissimilarity values below 0.8 were included in subsequent analyses.

**Figure S4.**
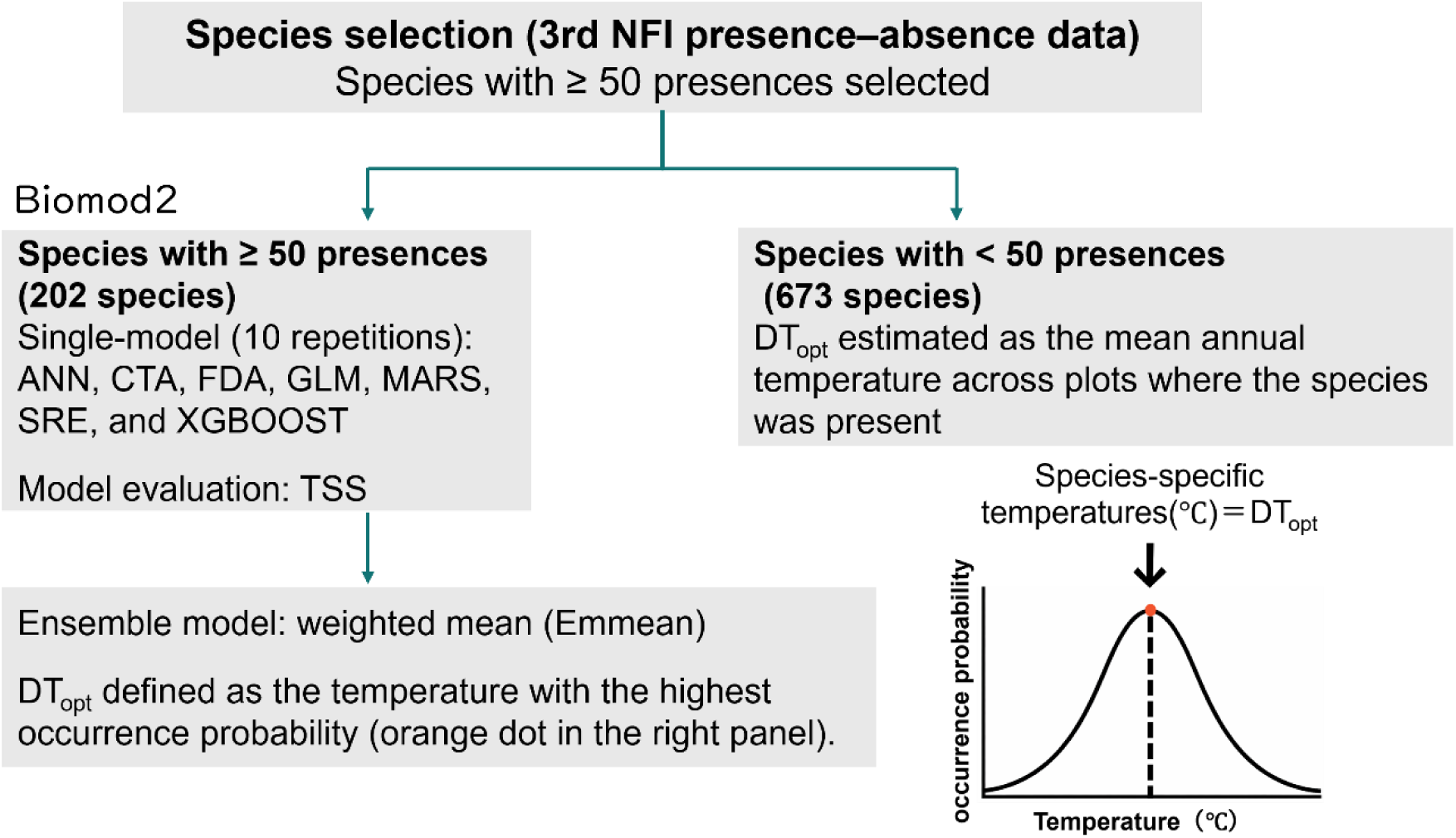
Flowchart of the modeling framework

**Figure S5.**
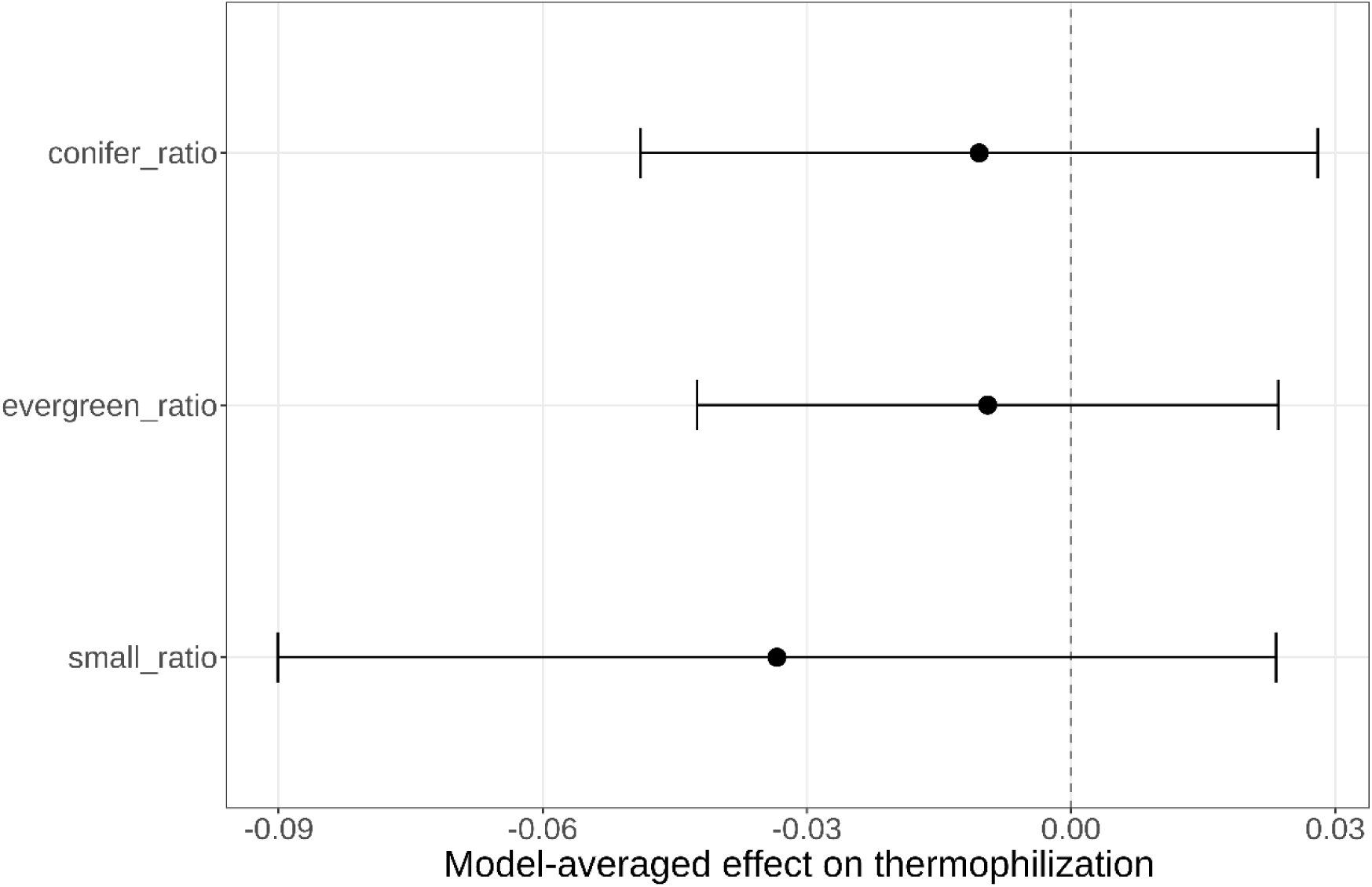
Model averaged coefficients of stand structure variables. The dots represent the estimated coefficients, and the error bars indicate the 95 percent confidence intervals. A variable is considered significant if its interval does not overlap with the dashed vertical line at zero.

**Figure S6.**
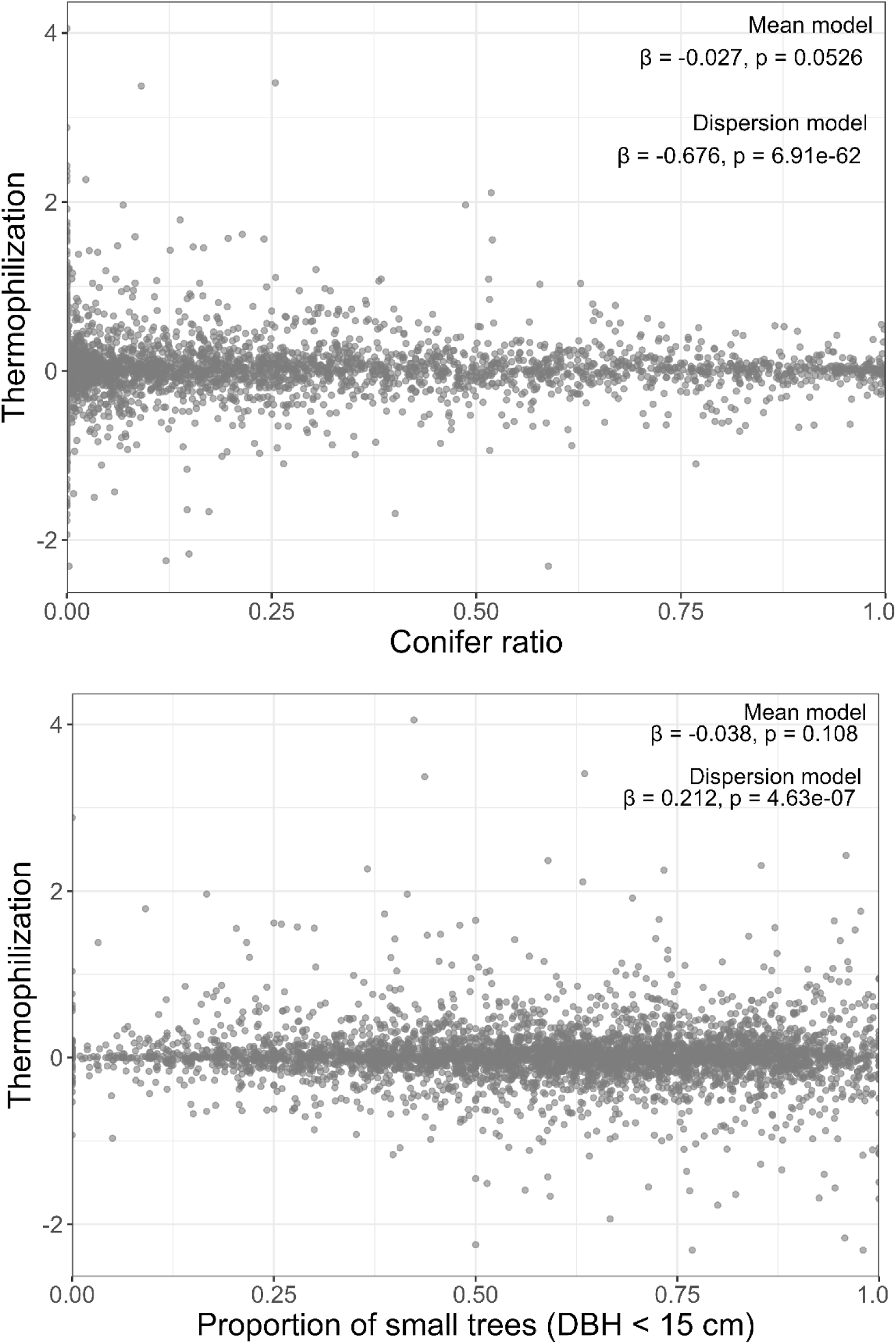
Relationship between the proportion of conifer ratio / small trees and thermophilization. Results are based on linear mixed-effects models (glmmTMB) with region included as a random factor. The model structure accounts for both mean and dispersion levels of thermophilization. Each point represents an individual study plot.

**Table S1.**
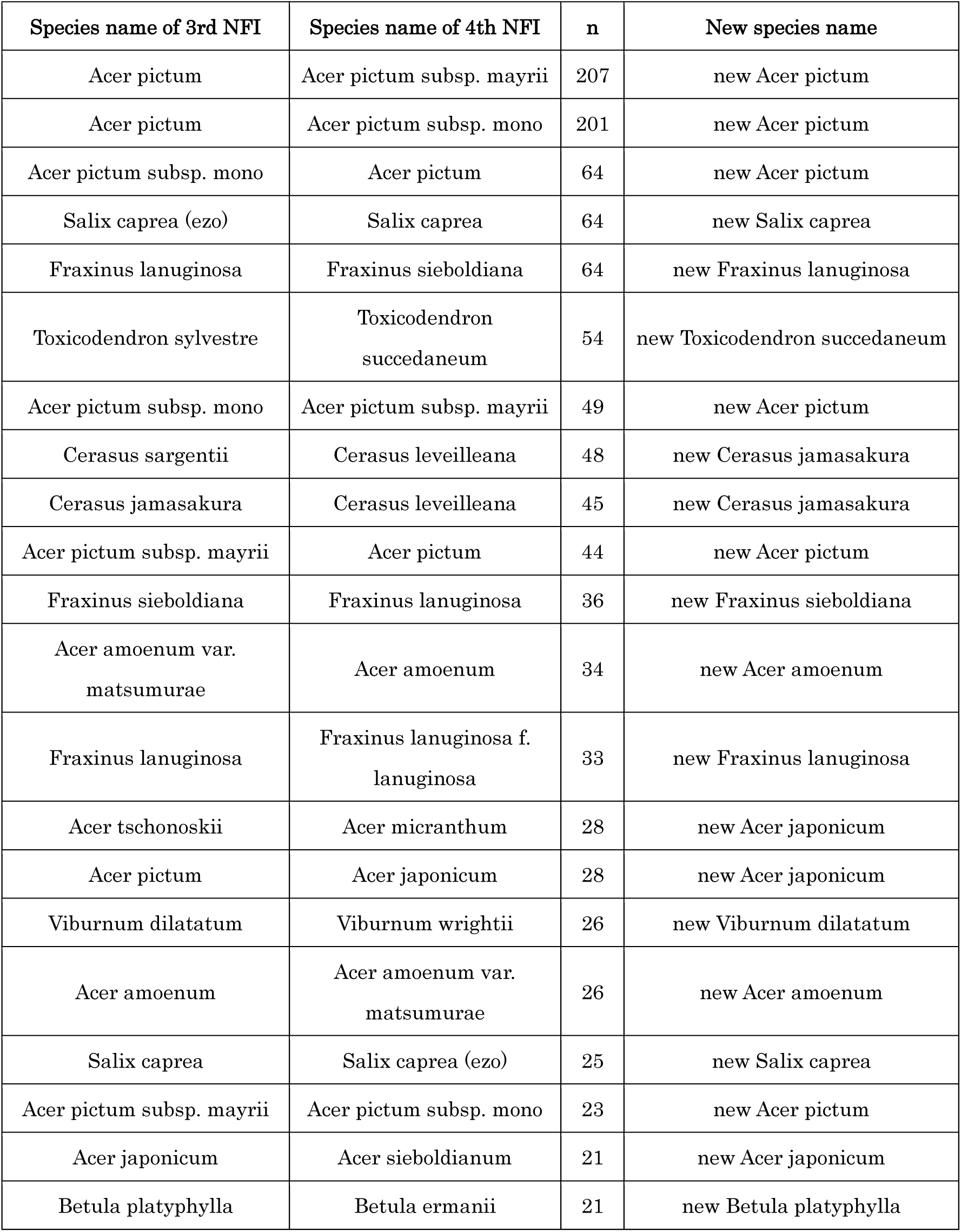
List of tree species newly defined for analysis. For each species, *n* indicates the number of plots in which the species was observed. Species that were clearly distinct and unlikely to be misidentified were excluded from redefinition.

| Species name of 3rd NFI | Species name of 4th NFI | n | New species name |
| --- | --- | --- | --- |
| Acer pictum | Acer pictum subsp. mayrii | 207 | new Acer pictum |
| Acer pictum | Acer pictum subsp. mono | 201 | new Acer pictum |
| Acer pictum subsp. mono | Acer pictum | 64 | new Acer pictum |
| Salix caprea (ezo) | Salix caprea | 64 | new Salix caprea |
| Fraxinus lanuginosa | Fraxinus sieboldiana | 64 | new Fraxinus lanuginosa |
| Toxicodendron sylvestre | Toxicodendron<br>succedaneum | 54 | new Toxicodendron succedaneum |
| Acer pictum subsp. mono | Acer pictum subsp. mayrii | 49 | new Acer pictum |
| Cerasus sargentii | Cerasus leveilleana | 48 | new Cerasus jamasakura |
| Cerasus jamasakura | Cerasus leveilleana | 45 | new Cerasus jamasakura |
| Acer pictum subsp. mayrii | Acer pictum | 44 | new Acer pictum |
| Fraxinus sieboldiana | Fraxinus lanuginosa | 36 | new Fraxinus sieboldiana |
| Acer amoenum var.<br>matsumurae | Acer amoenum | 34 | new Acer amoenum |
| Fraxinus lanuginosa | Fraxinus lanuginosa f.<br>lanuginosa | 33 | new Fraxinus lanuginosa |
| Acer tschonoskii | Acer micranthum | 28 | new Acer japonicum |
| Acer pictum | Acer japonicum | 28 | new Acer japonicum |
| Viburnum dilatatum | Viburnum wrightii | 26 | new Viburnum dilatatum |
| Acer amoenum | Acer amoenum var.<br>matsumurae | 26 | new Acer amoenum |
| Salix caprea | Salix caprea (ezo) | 25 | new Salix caprea |
| Acer pictum subsp. mayrii | Acer pictum subsp. mono | 23 | new Acer pictum |
| Acer japonicum | Acer sieboldianum | 21 | new Acer japonicum |
| Betula platyphylla | Betula ermanii | 21 | new Betula platyphylla |

**Table S2.** Regional distributions of thermophilization. The figure summarizes the number of plots (*n*) and the distribution of thermophilization values for each region. *Q1*, *mean*, and *Q3* denote the first quartile, mean, and third quartile, respectively.

| Region | n | Q1 | Median | Q3 |
| --- | --- | --- | --- | --- |
| Hokkaido | 1395 | -0.085 | 0.016 | 0.154 |
| Tohoku | 1089 | -0.103 | 0.005 | 0.110 |
| Kanto/Chubu | 587 | -0.102 | 0.008 | 0.122 |
| Kinki/Sanyo | 530 | -0.063 | 0.018 | 0.108 |
| Hokuriku/Sanin | 386 | -0.117 | -0.001 | 0.122 |
| Kyushu/Shikoku | 398 | -0.140 | 0.007 | 0.137 |
| Okinawa | 21 | -0.263 | -0.031 | 0.086 |

